# Insight into relationship between micro-consortia, nitrogen source and petroleum degradation at low temperature anaerobic condition

**DOI:** 10.1101/358838

**Authors:** Jicheng Yu, Chao Chen, Changjian Liu, Dongning Yu, Shuai Chen, Fenghao Yuan, Yang Fu, Qiu Liu

## Abstract

Biostimulation by addition nutrients has been proved to be an effective bioremediation strategies. Revealing response law of nitrogen source and structure characteristics of anaerobic petroleum degrading microorganisms microbial population will help us optimize nutrient to promote oil degradation. Anaerobic micro-consortia characteristics in the enrichment marine sediment samples with different nitrogen source, combining with analysis of the oil degradation rates were studied in this paper, as well as functional genes involved in petroleum degradation were also analyzed. On the basis of optimizing the best inorganic nitrogen sources and organic nitrogen sources, an effective medium was designed by response surface methodology that used for enriching petroleum degradation micro-consortia. Amplicon sequencing analysis showed that the population of microorganisms migrated obviously when enriched with different nitrogen sources. With the increase of oil degradation rate, the microbial diversity was significantly decreased, and concentrated on a limited number of genera. The reasonable proportions of *GammaProteobacteria, Bacteroidetes* and *Fusobacteria* made the greatest contribution to petroleum degradation. Metagenomic analysis unveiled the mixed nitrogen source promoted the expression of functional genes related to petroleum degradation such as the transfer of succinyl-CoA, synthesis of acetyl CoA and β-oxidation cycle, and was beneficial to degradation of petroleum at low temperature anaerobic condition.

**Originality Significance Statement:** Addition of nutrients can promote growth of indigenous petroleum degradation-related bacteria and be helpful to the rapid degradation of petroleum. Previous studies accurately characterized aerobic microorganisms on petroleum degradation. However, we still known little about anaerobic microorganisms in marine environment. Most biostimulation methods use inorganic salt as the main nutritional supplement to improve the efficiency of petroleum degradation, but effects of different nitrogen sources on diversity of microorganisms and distribution of functional genes related to petroleum degradation at anaerobic conditions are still unknown. In this research, the effects of nitrogen on petroleum biodegradation, anaerobic microconsortium structure and distribution of genes related to petroleum degradation were unveiled by using amplicon sequencing and metagenomic analysis.

## Introduction

Petroleum pollution frequently occurred in marine environment. Marine is such a special environment with high salt, low temperature and oligotrophy that removal of crude oil from the sea, especially from the seafloor, is much more difficult. Microorganisms act as one of the most important bio-degraders exhibit tenacious survival ability in the harsh marine environment (6), thus, bioremediation of oil contaminated seafloor is mainly dependent on marine indigenous microorganisms (13, 17). Previous studies accurately characterized aerobic microorganisms on petroleum degradation, meanwhile, the key degraders were well identified. However, we still known little about anaerobic microorganisms in marine environment.

Nowadays, many efforts have been made to explore suitable bioremediation strategies that can be applied to remove oil pollutants away from the seafloor (19, 39). Addition of nutrients and improving environmental conditions can promote growth of indigenous petroleum degradation-related bacteria and their petroleum degradation ability. So biostimulation by addition nutrients has been proved to be an effective bioremediation strategies for petroleum biodegradation (28, 30).

Because spilled crude oil brings additional carbon source into marine environment, which breaks the balance of nitrogen, in this case, the nitrogen for the indigenous microorganisms in marine environment are deficiency. Therefore, the supplement of nitrogen sources is undoubtedly important for promoting the rapid growth of marine indigenous microorganism, as well as accelerating the degradation rate of crude oil (23). Up to date, the relevant studies mainly focused on the feasibility of using inorganic nitrogen to facilitate the degradation of spilled oil at aerobic condition. Are organic nitrogen sources beneficial to the growth of indigenous petroleum degrading microorganisms? The nitrogen source response rule of petroleum degrading microorganisms in low temperature and anaerobic high salinity environment are still unknown. Composition of microorganisms that play a leading role in petroleum degradation also are still unknown.

In this research, the effects of nitrogen on petroleum biodegradation, anaerobic microconsortium structure and distribution of degradation genes were unveiled by using amplicon sequencing and metagenomic analysis. The aims of this study are to (i) Finding out the optimum combination of nitrogen nutrients for promoting petroleum degradation rate at low temperature anaerobic condition, (ii) revealing the involved microbial communities characteristics in the biostimulation enrichment samples with different nitrogen source by using amplicon sequencing, combining with analysis of the oil degradation rates, (iii) functional genes distribution involved in metabolic pathways of petroleum degradation by metagenomic analysis, further verification of the relationship between different nitrogen sources and oil degradation efficiency.

## Materials and Methods

### Description of sampling sites and process

The marine sediments were collected respectively from Xingang port of Dalian, China, which is the largest deepwater port in China. It is located on the coast of the Yellow Sea, which is near the Gulf of Bohai (Fig. 2A). Samples were collected 10–40 meters depth below the seawater, then put them in sterile valve bottles and then transported to the lab preserved at 4℃before the study. Temperatures of sampling sites were monitored to be 15-18ºC in October. All samples were numbered by sampling orders (11 sites such as E2, E4-7, E10-14 and E17). The mixture of the 4 samples (No. E5, E6, E13-14) was used for enrichment culture inoculum, other samples (No. E2, E4, E7, E10-12, E17) were as inoculants for evaluating the effect of petroleum degradation in this study (Fig. 2B).

### Enrichment medium of anaerobic microconsortiums

Oil degrading microorganisms were enriched in ASM medium (8) by adding crude oil as sole carbon source. The medium contained NaCl (30 g), MgSO_4_·7H_2_O (0.35 g), Na_2_HPO_4_(5 g), trace element solution (10 mL) and deionized water (1000 mL). The trace element solution was defined as 2 mg of CaCl_2_, 50 mg of FeCl_3_·6H_2_O, 0.5 mg of CuSO_4_, 0.5 mg of MnCl_2_ and 10 mg of ZnSO_4_·7H_2_O. The pH was adjusted to 7.5 before sterilization.

#### Culture conditions

Anaerobic culture approach was adopted by Li et al. (11). Briefly, brine bottles filled with ASM medium were put into an anaerobic glove box (DG250,Don Whitley Scientific Ltd, UK), vacuum pumped (66 kpa) three times, refilled with nitrogen gas (99.99%), followed by another three times of vacuum pumping (66 kpa) and refilling with mixed gases (H_2_ 10%, CO_2_ 5%, N_2_ 85%). Brine bottles were then sealed with butyl rubber plugs and static incubated at 15℃in dark. Anaerobic culture was carried out for 10 days.

### Anaerobic microconsortia enrichment treated with different nitrogen sources and evaluation of their degradation ability

Enrichment experiments were performed in ASM medium with crude oil (0.3%V V^-1^) as the sole carbon source, 2 g L^-1^ inorganic nitrogen sources such as NH_4_Cl, NaNO_3_, and NH_4_NO_3_ with organic nitrogen sources such as soybean flour, peanut meal flour, corn flour and bran were added as sole nitrogen source, respectively. 10 g of the mixture was inoculated into 50 mL medium in autoclave sterilized pressure culture bottle (250 mL). ASM medium with crude oil which worked as carbon source without any nitrogen source was used as control 1 for the evaluation of oil degradation rates, control 1 with mixture sediments inoculation was used as control 2 for the evaluation of the effect of nitrogen source on oil biodegradation in this research. All enriched samples were cultivated at 15℃under anaerobic condition for 10 days. The oil biodegradation rates were detected by UV spectrophotometry (Shimadzu UV-2450, Japan) at 225 nm, which were assessed as follows: η%=A_0_-A_1_/A_0_×100%η% means biodegradation rate; A_0_ means UV absorption of the control 1 extracted by petroleum ether; A_1_ means UV absorption of the enriched cultures extracted by petroleum ether (16). The cells in enriched cultures were collected for further amplicon sequencing and metagenomic analysis.

### Determination of optimum concentration of NH_4_NO_3_, Na_2_HPO_4_ and soybean flour for oil biodegradation

According to the oil degradation rates of above results, NH_4_NO_3_ was the best inorganic nitrogen source; and soybean flour was the best organic nitrogen source. Phosphorus source Na_2_HPO_4_ was an important factor for micro-consortia inoculation and oil biodegradation. Thus, concentrations of NH_4_NO_3_, soybean flour and Na_2_HPO_4_ were designed from 0.5 g L^-1^ to 3 g L^-1^, 1 g L^-1^ to 6 g L^-1^ and 0 g L^-1^ to 7 g L^-1^, respectively, to detect the optimum concentration of each single factor in ASM medium to obtain the higher oil biodegradation rate.

### Application of Response Surface Methodology and statistical design to the optimization of culture medium for better oil degradation

Based on the above results, three single factors (NH_4_NO_3_, soybean flour and Na_2_HPO_4_) at three levels [33] were applied and a series of 17 experiments (Table 1) were carried out according to the Box–Behnken design (BBD), and response surface methodology (RSM) was used to optimize the selected three significant variables(33), and the medium with the highest oil biodegradation rate was designated as YH was used for further experiments. The parameters and their levels were presented in Table 1. The statistical software package “Statistics Analysis System (SAS) 9.1” was used to analyze the experimental data. All variables were taken at a central coded value considered as zero. After the completion of experiments, oil degradation rate of each enrichment sample was evaluated. A multiple regression analysis of the data was performed for obtaining an empirical model which relates the response measured to the independent variables. Once the experiments were performed, the results were fitted with a second order polynomial equation:

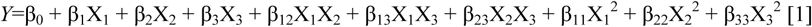

Where *Y* was measured response, β_0_ was the intercept term, β_1_, β_2_ and β_3_ were linear coefficients, β_12_, β_13_ and β_23_ were interaction coefficients, β_11_, β_22_ and β_33_ were squared coefficients, and X_1_, X_2_ and X_3_ were coded independent variables.

Statistical significance in the equation was determined by F-test. The coefficient of correlation (R^2^), adjusted coefficient of determination (R^2^ adj) and predicted coefficient of determination (R^2^ pred) were evaluated to investigate the model adequacies. The analysis of variance (ANOVA) was selected to test the statistical significance of the regression coefficients after selecting the most accurate model. Design-Expert Software was used to spot the response surface graphs. The optimum medium composition was verified through performing supplementary confirmation experiments at these conditions. The p-values of less than 0.05 meant statistically significant.

### DNA extraction and microbial diversity analysis of the samples

The enrichment cultures with higher petroleum degradation rate were selected as targets for microbial diversity analysis; therefore, the samples were enriched by the different nitrogen sources, NH_4_Cl, NaNO_3_, NH_4_NO_3_, soybean flour, and peanut meal flour (the enriched samples were designated as NCl, NaN, NN, DD and HS, respectively); in the meantime, samples enriched by YH medium was designated as YH. control 2 was used as control in this research to determine the microbial diversity, which was assessed by amplicon sequencing.

Total DNA extraction and 16S rRNA sequencing were performed by Novegene company (Beijing, China). Total DNA extraction was conducted with the FAST DNA^®^ Spin Kit for soil (MP Biomedicals, LLC, Solon,OH) according to the manufacturer’s instructions. DNA extracted from three technical replicates of each sample was pooled into one DNA sample to minimize any potential DNA extraction bias. OD value of the extracted DNA preparations is between 1.8~2.0.

DNA samples were amplified by PCR procedure using primer set F515 (5′-GTGCCAGCMGCCGCGG-3′) and R907 (5′ -CCGTCAATTCMTTTRAGTTT-3′) for the V4 region of the 16S rRNA gene (14). PCR was conducted using TransGen AP221-02 in a total volume of 20 μl with 4 μl 5×FastPfu Buffer, 2 μldNTPs (2.5 mM), 0.8μl of each primer (5 μM), 0.4 μl FastPfu polymerase, 0.2 μl BSA and 10ng of template DNA. PCR was performed in a GeneAmpR○ 9700(Applied Biosystems, U.S.) and the PCR conditions were as follows: 3 min at 95 °C; 27 cycles consisting of 30 s at 95°C, 30 seconds at 55°C and 45 seconds at 72°C; with a final extension step at 72°C for 10 min. The PCR products were purified using the UNIQ-10 PCR Purification Kit (Majorbio,Shanghai, China). After purification, the 16S rRNA V4 region PCR products were quantified using TBS-380 (Turner Biosystems USA). A mixture of the amplicons was sequenced on an Illumina MiSeq platform according to the standard protocols.

Each sample was sequenced for three technical replicates. The sequences were clustered into operational taxonomic units (OTUs) by setting a 0.03 distance limit (equivalent to 97% similarity) by using the MOTHUR program (27). From the cluster file, OTU richness indices such as Chao and abundance-based coverage (ACE) estimators, Shannon diversity index and the Good’s coverage were determined by the Mothur program based on observed OTUs defined at 97% sequence identity for each sample. Sequences were also phylogenetically assigned to taxonomic classifications by using an RDP classifier Bayesian Algorithm(35). The relative abundance of a given phylogenetic group was the sequence number of the affiliated group divided by the total number of sequences per sample.

### Metagenomic sequencing

Isolation, purification and detection of metagenomic DNA was the same as the above described. The samples which were named CK, NN, DD and YH were selected as targets for metagenomic sequencing, respectively. Sequencing libraries were generated using NEBNext^®^ Ultra™ DNA Library Prep Kit for Illumina (NEB, USA) following manufacturer’s recommendations and index codes were added to attribute sequences to each sample. The DNA fragments were sequenced on an IlluminaHiSeq platform and paired-end reads were generated. Preprocessing the Raw Data obtained from the IlluminaHiSeq sequencing platform using Readfq V8, https://github.com/cjfields/readfq was conducted to acquire the Clean Data for subsequent analysis. ForSingle sample assembly, MEGAHIT software (v1.0.4-beta) was used to assemble the Clean Data. All samples’ Clean Data are compared to each Scaffolds respectively by SoapAligner software (soap 2.21) to acquire the PEreads were not used. Furthermore, all the reads which were not used in the forward step of all samples are combined and then use the software of SOAPdenovo (V2.04) / MEGAHIT (v1.0.4 -beta) for mixed assembly. The Scaftigs (≥ 500 bp) assembled from both single and mixed are all predicted the ORF by MetaGeneMark (V2.10, http://topaz.gatech.edu/GeneMark/) software. For ORF predicted, CD-HIT software (V4.5.8, http://www.bioinformatics.org/cd-hit) is adopted to redundancy and obtain the unique initial gene catalogue. The Clean Data of each sample was mapped to initial gene catalogue using SoapAligner (soap2.21) and get the number of reads to which genes mapped in each sample. Filter the gene which the number of reads less or equal to 2 in each sample and obtain the gene catalogue (Unigenes) eventually used for subsequently analysis.

### Bioinformatic analysis

DIAMOND software (V0.7.9, https://github.com/bbuchfink/diamond/) is used to blast the Unigenes to the sequences of Bacteria, Fungi, Archaea and Viruses which are all extracted from the NR database (Version: 20161115, https://www.ncbi.nlm.nih.gov/) of NCBI. Adopt DIAMOND software (V0.7.9) to blast Unigenes to functional database with the parameter setting of blastp, -e 1e-5 (12). Functional database excludes KEGG database (Version 201609, http://www.kegg.jp/kegg/), eggNOG database (Version 4.5, http://eggnogdb.embl.de/#/app/home), CAZy database (Version 20150704, http://www.cazy.org/). For each sequence’s blast result, the best Blast Hit is used for subsequent analysis(12). Statistic of the relative abundance of different functional hierarchy, the relative abundance of each functional hierarchy equal the sum of relative abundance annotated to that functional level.

Metagenomic data was also analyzed with standalone BLASTX v2.2. The key functional genes involving the petroleum degradation at anaerobic condition in this research and subsequently were annotated with MEGAN5 (reference: Improved metagenome analysis using MEGAN5).

## Results

### Effects of nitrogen sources and concentration of three single factors on oil biodegradation

Nitrogen source is important for the cycle of microorganisms’ life, hence in this research, inorganic and organic nitrogen sources, NH_4_Cl, NaNO_3_, NH_4_NO_3_, and soybean flour, peanut meal flour, corn flour, bran were used to identify their roles on oil biodegradation. In the studied inorganic nitrogen sources, NH_4_NO_3_ had better effect on oil biodegradation, the degradation rate was up to 44.94%, others were 38.47% (NH_4_Cl) and 36.65% (NaNO_3_), respectively (Fig. 1A); moreover, compared to inorganic nitrogen sources, organic nitrogen sources were more adaptable for oil biodegradation. The addition of soybean powder was the most conducive to oil biodegradation, the degradation rate was 62.61%, and others were 56.52% (peanut meal flour), 49.15% (corn flour) and 44.73% (bran), respectively.(Fig. 1A).

The above results indicated that organic nitrogen source was better than inorganic nitrogen source on oil biodegradation. In this case, in order to design an optimum medium, the best inorganic nitrogen source NH_4_NO_3_ and the best organic nitrogen source soybean flour were selected as ingredients to determine their best concentration by single factor experiment. Phosphorus was also important for oil biodegradation; thus, its concentration was designed by single factor experiment. When added 2 g L^-1^ NH_4_NO_3_, 2 g L^-1^ soybean flour and 5 g L^-1^ Na_2_HPO_4_ in ASM medium(Fig. 1B–D), the oil biodegradation rates of micro-consortia were higher. Consequently, further we use response surface methodology to design the optimum medium according to the results obtained in this part.

**Figure 1.**
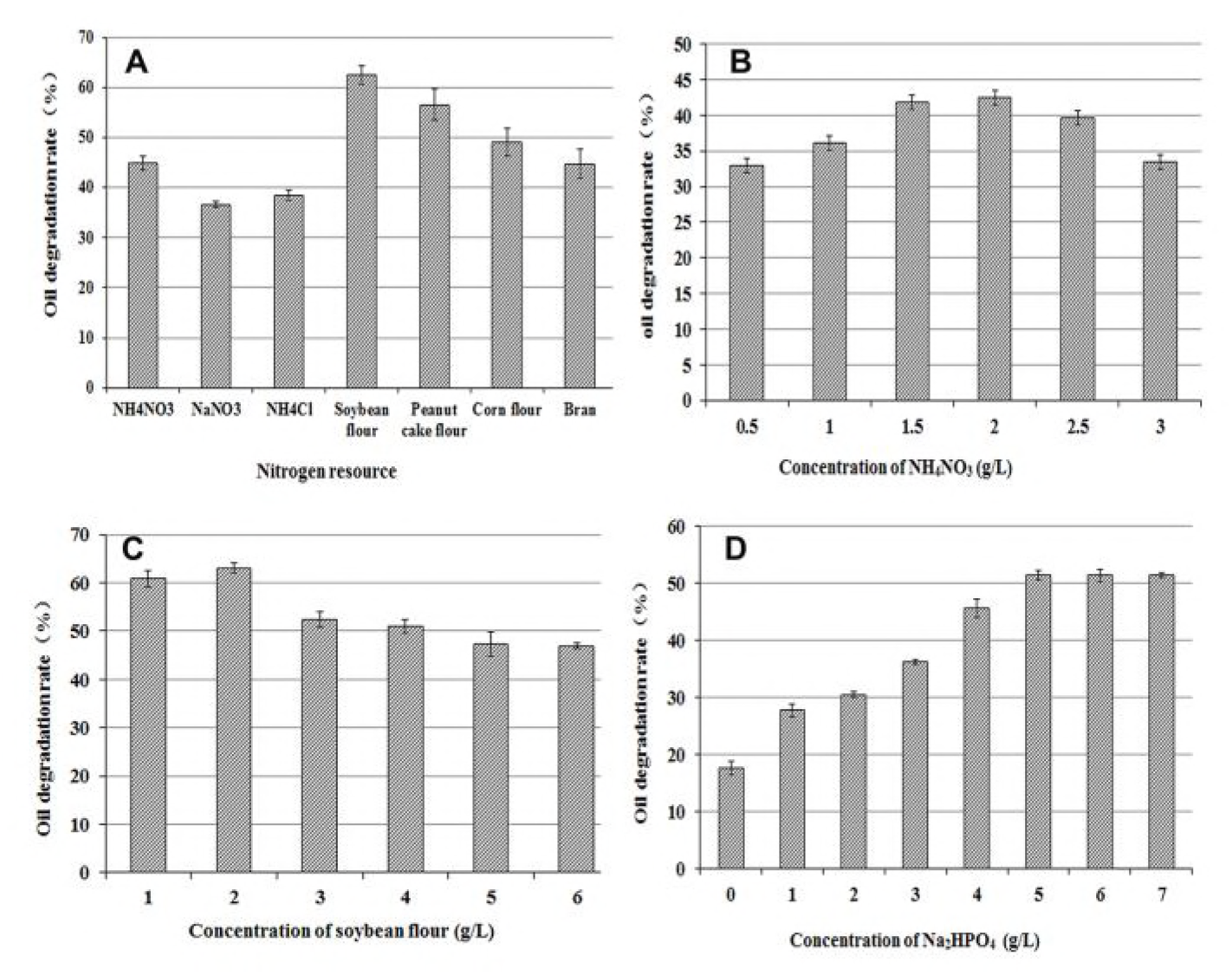
Effects of nitrogen sources and phosphate on oil degradation performance of microorganisms. (A) Effects of different nitrogen source on oil degradation performance. B and C Effects of concentration of NH_4_NO_3_, and soybean powder on oil degradation performance. (D) Effects of concentration of Na_2_HPO_4_ on oil degradation performance

### Optimum medium designing by response surface methodology and statistical analysis

A Box-Behnken design was applied to investigate the interactive effects of NH_4_NO_3_, soybean flour and Na_2_HPO_4_ on oil biodegradation. 17 experiments were performed at different levels of three factors (Table 1). The results were analyzed by SAS ANOVA procedure (Table 2). The second order polynomial equation for microbial oil degradation rate obtained from RMS was:

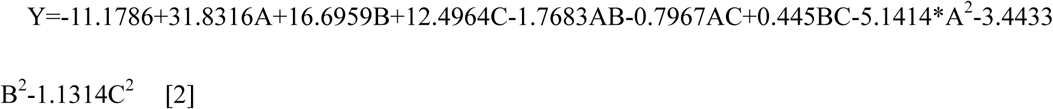

In which Y is the oil degradation rate of microorganisms, A, B and C are concentrations of soybean flour, NH_4_NO_3_ and Na_2_HPO_4_, respectively.

**Table 1.**
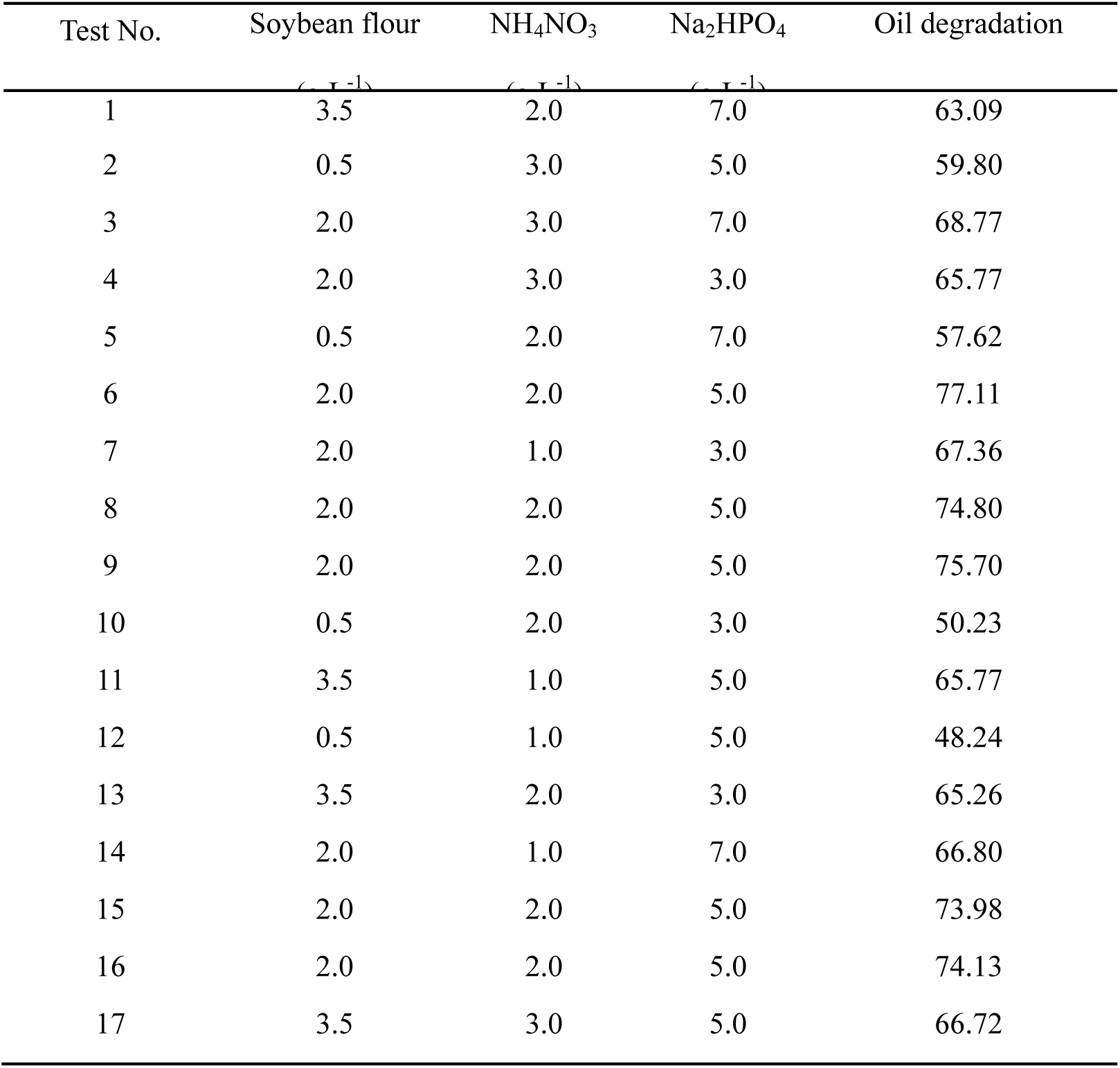
Box–Behnken design and results. The test NO. 2 showed the highest oil degradation rate.

| Test No. | Soybean flour | NH<sub>4</sub>NO<sub>3</sub> | Na<sub>2</sub>HPO<sub>4</sub> | Oil degradation |
| --- | --- | --- | --- | --- |
| 1 | 3.5 | 2.0 | 7.0 | 63.09 |
| 2 | 0.5 | 3.0 | 5.0 | 59.80 |
| 3 | 2.0 | 3.0 | 7.0 | 68.77 |
| 4 | 2.0 | 3.0 | 3.0 | 65.77 |
| 5 | 0.5 | 2.0 | 7.0 | 57.62 |
| 6 | 2.0 | 2.0 | 5.0 | 77.11 |
| 7 | 2.0 | 1.0 | 3.0 | 67.36 |
| 8 | 2.0 | 2.0 | 5.0 | 74.80 |
| 9 | 2.0 | 2.0 | 5.0 | 75.70 |
| 10 | 0.5 | 2.0 | 3.0 | 50.23 |
| 11 | 3.5 | 1.0 | 5.0 | 65.77 |
| 12 | 0.5 | 1.0 | 5.0 | 48.24 |
| 13 | 3.5 | 2.0 | 3.0 | 65.26 |
| 14 | 2.0 | 1.0 | 7.0 | 66.80 |
| 15 | 2.0 | 2.0 | 5.0 | 73.98 |
| 16 | 2.0 | 2.0 | 5.0 | 74.13 |
| 17 | 3.5 | 3.0 | 5.0 | 66.72 |

The ANOVA of the quadratic regression model demonstrated that Eq. [2] is a significant model, which is evident that it is from the F-test with a low probability value (Table 2). Values of “Prob. > F” less than 0.05 indicated that model term was significant, and “Prob. > F” less than 0.01 indicated that model term was very significant. In the present work, The effects of A, the effects of interaction between A and B,A and C, and the effects of the square effects of A, B, C were significant for oil degradation ability of microorganisms. Thus, it was further proved that nitrogen sources, especially organic nitrogen sources (A and A^2^ was very significant), were crucial for microbial oil degradation. The coefficient of determination (R^2^) for oil degradation ability of microorganisms was calculated as 0.9750, which indicated that 97.50% of the total variability in the response could be explained by this model. The present R^2^-value reflected an acceptable fit between the experimentally observed and predicted values. Therefore, the model could be used to predict the oil degradation ability of microorganisms within the limits of the experimental factors.

**Table 2.**
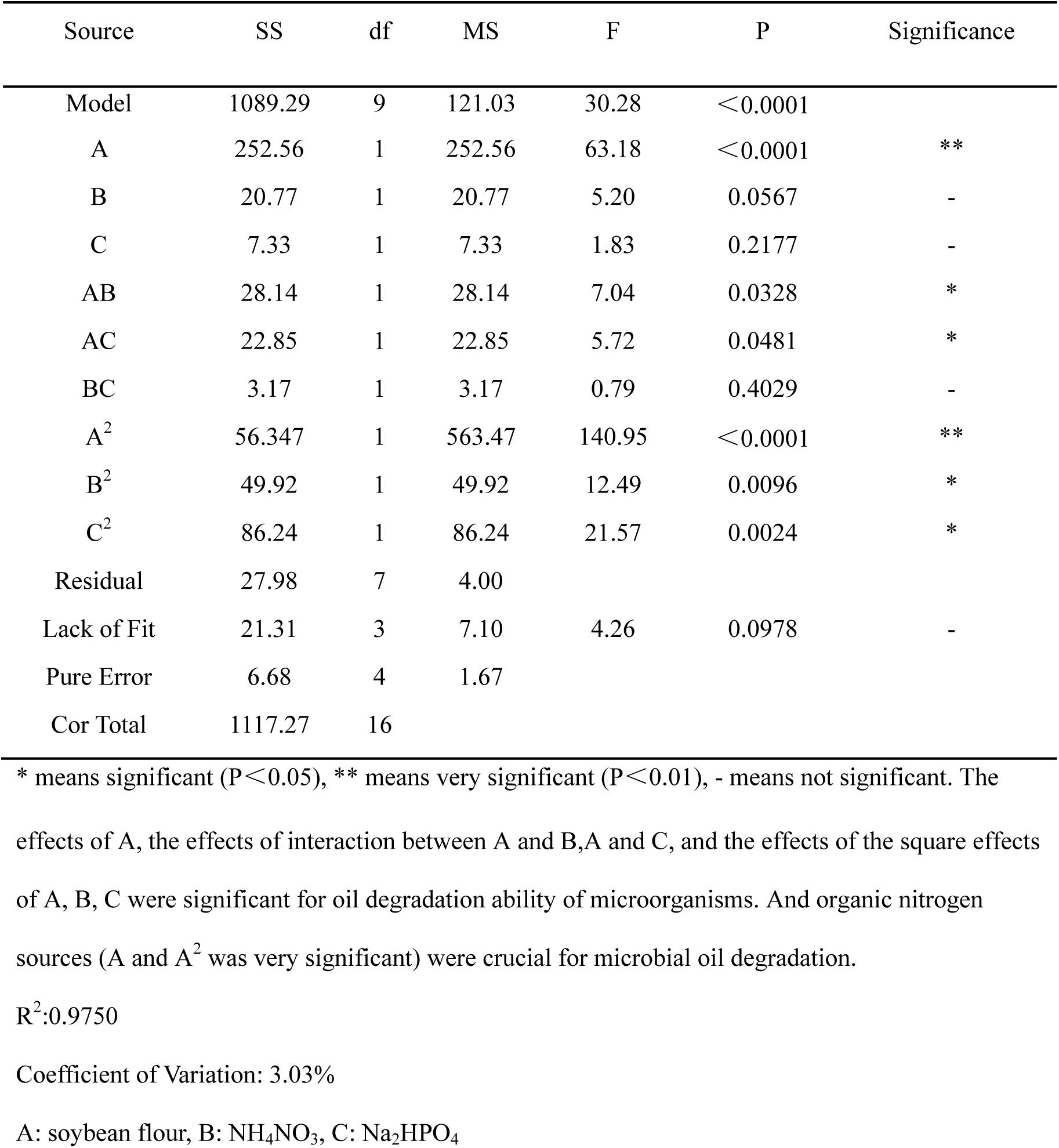
The analysis of variance of Box-Behnken design results. The effects of A, the effects of interaction between A and B,A and C, and the effects of the square effects of A, B, C were significant for oil degradation ability of microorganisms. And organic nitrogen sources (A and A^2^ was very significant) were crucial for microbial oil degradation.

Optimum conditions for the maximum oil degradation were determined by response surface analysis and also estimated by regression equation. The optimum medium were soybean flour 2.33 g L^-1^, NH_4_NO_3_2.16 g L^-1^, and Na_2_HPO_4_ 5.13 g L^-1^, the predicted oil degradation rate were 75.91%. We investigated the accuracy of the model by carrying out the batch experiment under optimal operation conditions. Seven repeated experiments were performed. The average value of oil degradation rate (74.93±0.84%) was deeply close to the response predicted (75.91%) by the regression model.

### Evaluation of universality of the optimized medium YH

In order to evaluate the universality of the optimized medium, seven marine sediments (Fig. 2B, sampling orders were No. E2, E4, E7, E10-12, E17 respectively) with different petroleum contents were inoculated in the optimized medium and cultured under the same condition to detect the oil biodegradation rates, the results showed that the oil biodegradation rates were between 71.86% – 86.61% (Fig. 2C), which verified the universality of the optimized medium(the sample number was YH enriched with the optimized medium).

**Figure 2.**
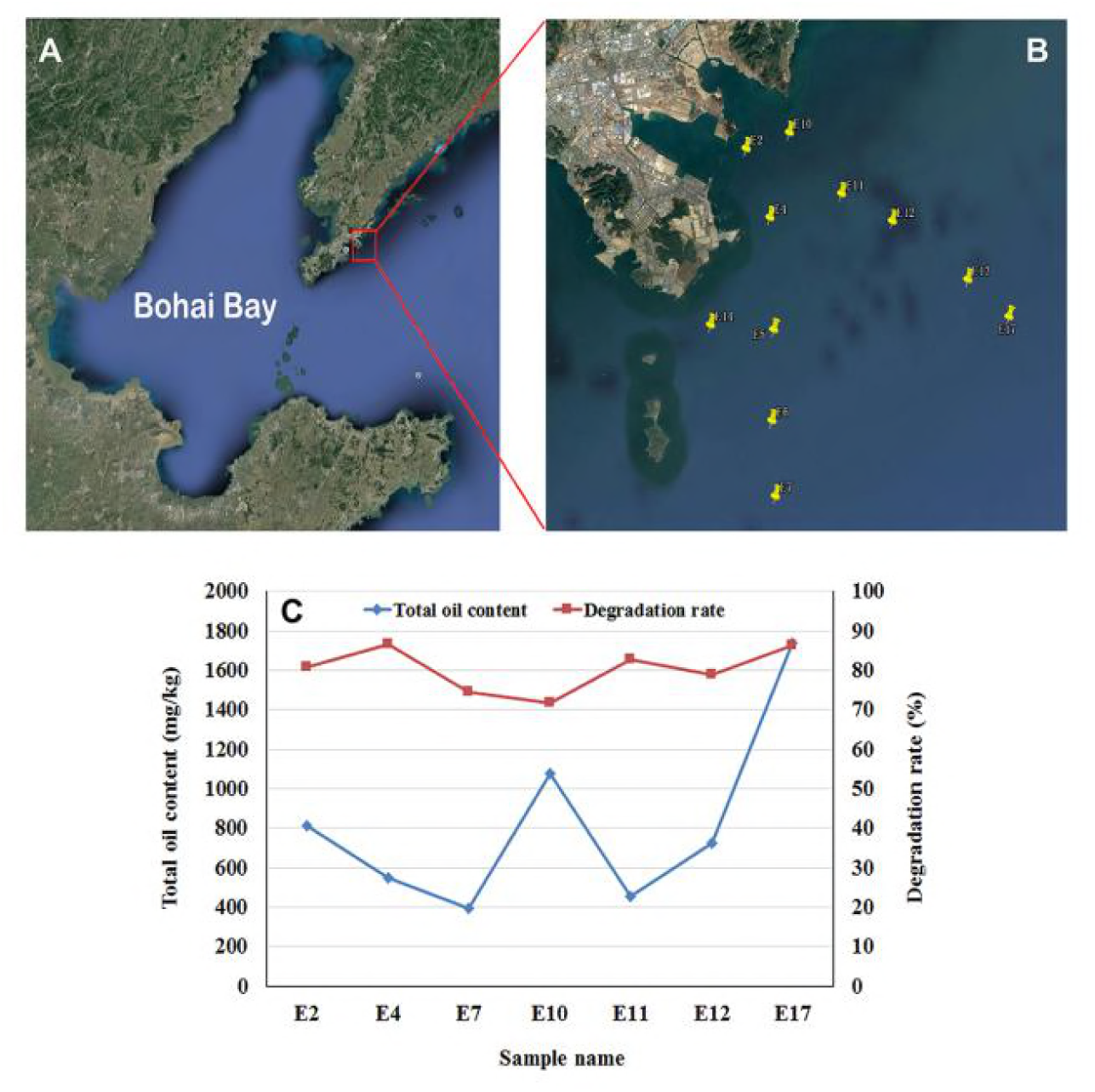
Map of sample collection locations and evaluation of petroleum degradation performance of samples enriched with optimized medium (A) Map of sample collection locations, (B) All samples were numbered by sampling orders. (C) Evaluation of petroleum degradation performance of samples enriched with optimized medium.

### Effects of nitrogen source on bacterial diversity

Marine sediments that enriched by different nitrogen sources were exhibited different bacteria diversities, with different bacteria abundance, which showed different petroleum biodegradation abilities (Table 3). OTU number in control 2 (CK) without any nitrogen source was the highest (414), and in the enriched ones, OUT numbers in samples enriched by organic nitrogen sources were lower than that enriched in inorganic nitrogen sources, But oil biodegradation rates indicated that micro-consortia in enriched samples can better consume petroleum as carbon source than CK, and organic nitrogen sources was better than inorganic nitrogen sources. Sample YH, with mixture of organic and inorganic nitrogen sources in it, showed the highest oil biodegradation rate, which indicated that different kinds of nitrogen sources can enrich different type of biodegradable microorganism groups.

**Table 3.** Analysis of abundance and diversity of microorganisms in different treatment samples with different nitrogen source. Marine sediments that enriched by different nitrogen sources were exhibited different bacteria diversities (Shannon index), with different bacteria abundance (Simpson index), which showed different petroleum biodegradation abilities. Note that *seven samples.CK, DD, HS, NCl, NN, NaN and YH were the sample enriched in ASM medium without nitrogen source addition, with 2 g L^-1^ soybean flour addition, with 2 g L^-1^ peanut cake flour addition, with 2 g L^-1^ NH_4_Cl addition, with 2 g L^-1^ NH_4_NO_3_ addition, with 2 g L^-1^ NaNO_3_ addition, with the optimized mixture nitrogen source addition, respectively.

| Sample No.* | Reads | OTUs | ACE | Chao | Coverage rate | Shannon index | Simpson index | Oil degradation rate(%) |
| --- | --- | --- | --- | --- | --- | --- | --- | --- |
| CK | 14067 | 414 | 435<sup>#</sup> | 437 | 0.996730 | 4.09 | 0.0559 | 39.26±1.36 |
| NaN | 30625 | 370 | 402 | 401 | 0.998171 | 2.73 | 0.148 | 39.77±2.71 |
| NCl | 19229 | 376 | 406 | 398 | 0.997192 | 3.3 | 0.1037 | 42.95±0.21 |
| NN | 21554 | 399 | 413 | 410 | 0.998470 | 3.31 | 0.1431 | 49.66±0.86 |
| HS | 24214 | 74 | 130 | 95 | 0.998968 | 1.44 | 0.4672 | 64.93±0.69 |
| DD | 32853 | 150 | 203 | 203 | 0.998356 | 2.33 | 0.1613 | 66.44±0.33 |
| YH | 29463 | 105 | 206 | 194 | 0.997167 | 1.94 | 0.2154 | 73.83±0.84 |

medium without nitrogen source addition, with 2g L<sup>-1</sup> soybean flour addition, with 2g L<sup>-1</sup> peanut
cake flour addition, with 2g L<sup>-1</sup> NH<sub>4</sub>Cl addition, with 2g L<sup>-1</sup> NH<sub>4</sub>NO<sub>3</sub> addition, with 2g L<sup>-1</sup>

Bacterial communities in enriched samples were identified at phylum and genus level. And distributions of microbes in the samples enriched with organic nitrogen sources were obviously different from that in samples enriched with inorganic nitrogen sources. Top 8 phyla (relative abundance >1% in at least one enrichment sample), *Proteobacteria, Firmicutes, Cyanobacteria, Bacteroidetes, Acidobacteria, Planctomycetes, Fusobacteria* and *Chloroflexi* were analyzed (Fig. 3A). In CK, *Proteobacteria* (57.3%) was the most predominant, which was followed by *Firmicutes*(13.8%) *Cyanobacteria*(11.2%) a*nd Bacteroidetes*(6.5%), the total abundance of these four phyla was up to 88.8%. Previous study showed that the addition of organic matter to crude-oil amended sediment microcosms significantly increased the mineralization rates for hydrocarbons and particularly enriched groups of *Proteobacteria* (20, 24). Furthermore, in this research, we found that the addition of inorganic nitrogen sources were also enriched particular groups of *Proteobacteria,* but with the increase of oil degradation rates, its relative abundance gradually decreased. In enrichment sample NN, the total abundance of four phyla with the highest relative abundance were up to 91.6%, they were *Proteobacteria* (59.8%), *Firmicutes* (24.3%), *Cyanobacteria* (3.6%) and *Bacteroidetes*(3.9%); meanwhile, the oil biodegradation rate was up to 49.66%. The composition of the top four phyla in CK with lower oil biodegradation rate (39.26%) were the same as that in NN, the ratio of *Firmicutes*in in the NN was higher than that in CK, which indicated that *Firmicutes* might be a better petroleum degradation groups. In enrichment samples NaN and NCl, *Fusobacteria* and *Firmicutes* were the second and third predominant phyla, the relative abundance of the top four phyla were 94.5% and 90%, respectively, and the oil biodegradation rates were 39.77% and 42.95%, respectively, which were lower than that in enriched sample NN. The results above indicated that *Firmicutes* was a good degrader when using inorganic nitrogen source as sole nitrogen source.

**Figure 3.**
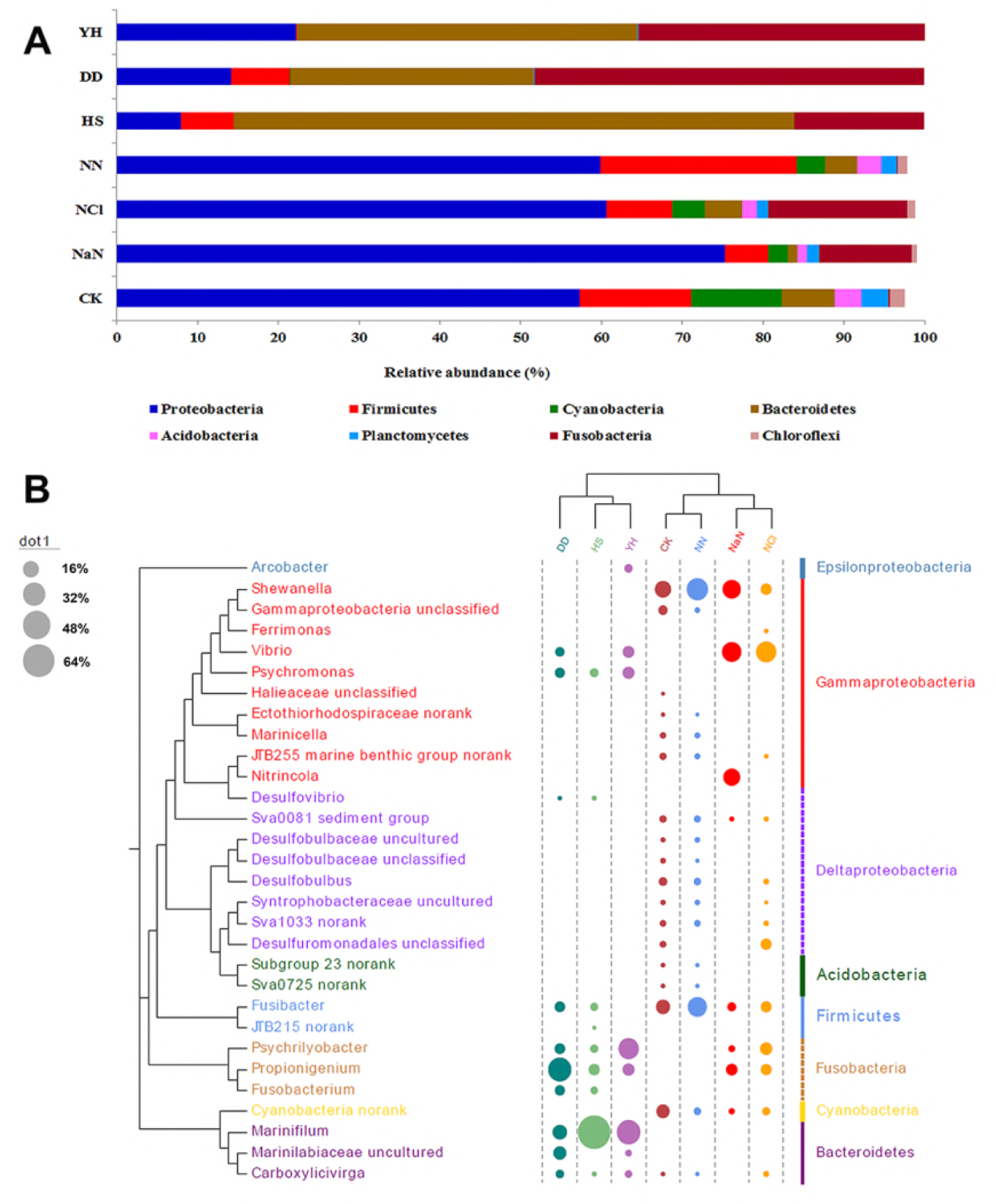
Phylum level (A) and genus level (B) distribution of bacterial community in the different nitrogen treatments. Note that NCl, NaN, NN, DD, and HS means the samples were enriched with the different nitrogen sources such as NH_4_Cl, NaNO_3_, NH_4_NO_3_, soybean flour, and peanut meal flour. YH means the samples enriched with YH medium was designated by the response surface experiment. CK means ASM medium control with crude oil which worked as carbon source without any nitrogen source, but with mixture sediments inoculation

In organic nitrogen sources enriched samples DD, HS and YH, the oil degradation rates were all higher than that in the samples with inorganic nitrogen sources, *Proteobacteria, Bacteroidetes* and *Fusobacteria* were the most dominant phyla, and the ratio of *Proteobacteria* was lower than that of *Bacteroidetes* and *Fusobacteria*; hence, the petroleum degradation rates of sample DD, HS and YH were 66.44%, 64.93% and 73.83, respectively, and compared to the diversities and ratios of microbes in sample DD, HS and YH, we can identified that these three top phyla were all dominant in oil degradation, but the ratio was more important. With the increase of degradation rate, the abundance ratio of the three dominant phylum was closer and more coordinated in sample YH, *Proteobacteria, Bacteroidetes* and *Fusobacteria* which were accounted for 22.16%, 42.11%, and 35.43%, respectively, the total ratio of which were up to 99.7%.

At the genus level, we compared the major genus (relative abundance >1% in at least one enrichment sample) which belong to 6 phyla such as *Proteobacteria, Firmicutes, Cyanobacteria, Bacteroidetes, Acidobacteria* and *Fusobacteria* (Fig. 3B). According to cluster analysis, the microbial community in sample CK was similar to that in the enrichment samples with inorganic nitrogen sources, and the microbial community in enrichment sample YH was similar to that in enrichment samples with organic nitrogen sources.

The sample CK involved more genera (18 genus with relative abundance >1%) than others, and among the 18 genus, *Shewanella* was dominant, accounting for 16.8%, followed by *Fusibacter* (12.6%) and *Cyanobacteria*(11.2%), and total abundance of the top 18 genera was up to 75.6%. With inorganic nitrogen sources enrichment, the number of the genus which were greater than 1% relative abundance were 8, 14, and 16 in the samples enriched by NaNO_3_ (NaN), NH_4_Cl (NCl) and NH_4_NO_3_(NN), respectively. And total abundance of the genera with 1% relative abundance was up to 84.25%, 82.07% and 81.43%, respectively. the microbial community in the enrichment samples with inorganic nitrogen sources was similar to that in sample CK, the diversity of microorganism is concentrated in *Proteobacteria*, and the microbial groups related to petroleum degradation are concentrated in *Gammaproteobacteria*.

There were 8, 9 and 10 genera with relative abundance which were greater than 1% in samples YH, HS and DD, respectively. In sample YH, the highest proportion of genus was *Marinifilum*, followed by *Psychrilyobacter*, the ratios of *Propionigenium, Psychromonas*, and *Vibrio* were basically the same, which were 8.89%, 8.86%, and 8.41%, respectively. And the total abundance of the genera with 1% relative abundance was up to 96.48%, 97.09% and 99.18%, respectively. the microbial community in enrichment samples with organic nitrogen sources was similar to that in enrichment sample YH, the diversity of microorganism is not concentrated in *Proteobacteria*, but is reflected in the diversity of phyla, and the microbial groups related to petroleum degradation are concentrated on *Gammaproteobacteria, Fusibacteria* and *Bacteroidetes*.

In enrichment sample YH, the 5 genera with the highest relative abundance were *Marinifilum, Psychrilyobacter, Propionigenium, Psychromonas*, and *Vibrio*. At anaerobic conditions, these five genera were the main microbial group for petroleum degradation, which account for 88.48% in the sample YH; however, which kind of collaboration between these microorganisms in the process of oil degradation was not clear. This result was consistent with the result at phylum level, that is, with the increase of oil degradation rate, the microbial diversity was significantly decreased, and concentrated on a limited number of genera. For example, in sample YH with the highest oil degradation rate, only 8 dominant genera were found, which accounting for 99.2% of the total biomass. The 8 dominant genera were *Marinifilum* (relative abundance 35.7%), *Psychrilyobacter* (26.6%), *Psychromonas*(8.9%), *Propionigenium* (8.9%), *Vibrio* (8.4%), *Arcobacter* (4.4%), *Carboxylicivirga* (3.4%) and *Marinilabiaceae* uncultured (2.9%). The genus with the highest relative abundance also changed from *Shewanella* (16.8%) in the sample CK to *Marinifilum*(35.7%) in sample YH. The results showed that the population of microorganisms migrated obviously when enriched with different nitrogen sources.

### Metagenomic sequencing and the basic information analysis

The enrichment samples CK, NN, DD and YH were used as an object for the analysis of the macrogenome. After redundancy, we obtained 25023.23Mbp clean data, these data were used to assemble and obtain 218979656 bp of Scaftigs with SOAP denovo software, and at last 228151 ORFs were obtained. The basic information in the four enrichment samples named NN, DD, YH and CK was shown in the petal graph (Fig. 4). The core value (23935) indicated the number of genes that shared in the four enrichment samples, which account for 10.49% of the total ORFs, the proportion of the shared genes was low. The numerical value in the petals indicated the difference between the gene number in each sample and the shared gene number from each sample. The values in parentheses indicated the number of genes and the number of unique genes in each enrichment sample (Fig. 4). The results showed that specific genes were obviously different from each other after enriched by different nitrogen sources, and there should be unique genes in each sample, including genes that might be related to petroleum degradation.

**Figure 4.**
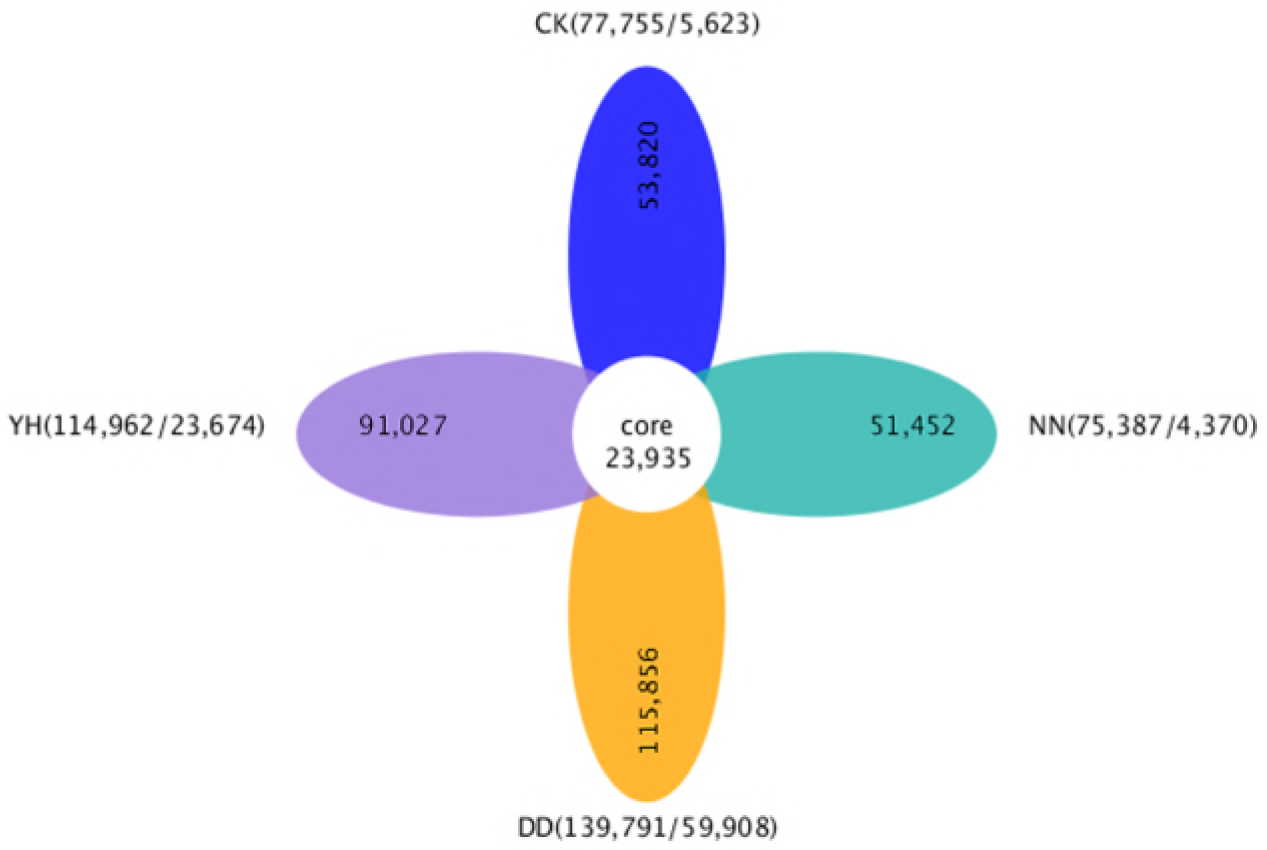
The basic information in the four enrichment samples named NN, DD, YH and CK. The core value (23935) in the petal graph indicated the number of genes that shared in the four enrichment samples. The numerical value in the petals indicated the difference between the gene number in each sample and the shared gene number from each sample. The values in parentheses indicated the number of genes and the number of unique genes in each enrichment sample.

### Analysis of the key genes of micro-consortia involved in anaerobic petroleum degradation by metagenomic sequencing

The enzymes involved in alkanes degradation pathway in anaerobic conditions such as alkylsuccinate/ benzosuccinates syhthase (ASS/BSS), succinyl-CoA: acetate CoA-transferase (SAcT), succinate dehydrogenase/fumarate reductase (SD/FR), acelyl-CoA carboxylase carboxyl transferase(ACCT) and acetyl-CoA synthetase (ACS). It is known from the experimental results that these key gene copies were different in four enrichment samples by metagenomic analysis. The organic nitrogen sources (sample DD) was the most beneficial to increase the number of the genes encoding ASS, BSS, ACCT, and SD/FR enzymes, especially for the ass and bss genes, there were 96 copies of ass gene and 89 copies of bss gene respectively, the mixture nitrogen (sample YH) source was followed. In addition, the mixed nitrogen promoted the transfer of succinyl-CoA, synthesis of acetyl CoA. For the copies of the genes encoding succinyl-CoA: acetate CoA-transferase and acetyl-CoA synthetase were the most,and they were 11 and 22 in the YH samples respectively (Fig. 5B).

**Figure 5.**
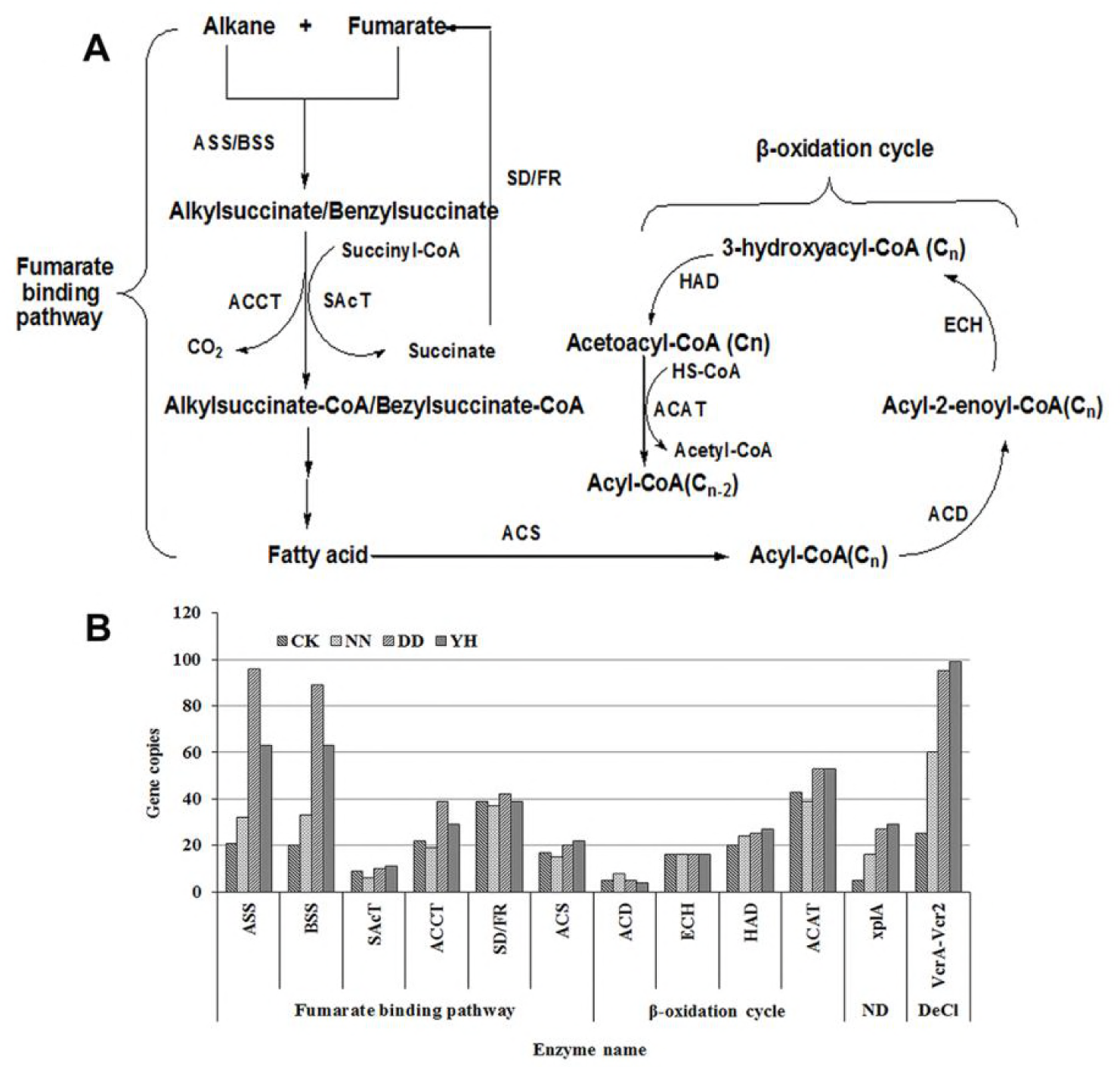
Proposed pathway of anaerobic degradation of alkanes(A) and the copy number difference of the genes involved anaerobic degradation of petroleum in four enrichment samples(B). Note that **Fumarate biding pathway involves the enzymes:** ASS Alkylsuccinate syhthase, BSS Benzosuccinates syhthase, SAcT Succinyl-CoA:acetate CoA-transferase, ACCT acelyl-CoA carboxylase carboxyl transferase, SD/FR Succinate dehydrogenase/fumarate reductase, ACS Acetyl-CoA synthetase. **β-oxidation cycle involves the enzymes:** ACD acyl-CoA dehydrogenase, ECH enoyl-CoA hydratase, HAD 3-hydroxyacyl-CoA dehydrogenase/ 3-hydroxybutyryl-CoA dehydrogenase, ACAT acetyl-CoA acetyltransferase. **ND and DeCl:**The pathway of nitramine degradation and Dechlorination of compound involves the key enzymes: xplA nitric oxide synthase, VcrA-Vcr2 reductive dehalogenase

In both aerobic and anaerobic conditions, microorganisms degrade petroleum must involve β-oxidation cycle (7, 25). We also analyzed the complete β-oxidation pathway involving the key enzymes acyl-CoA dehydrogenase(ACD), enoyl-CoA hydratase(ECH), 3-hydroxyacyl-CoA dehydrogenase/3-hydroxybutyryl-CoA dehydrogenase(HAD) and acetyl-CoA acetyltransferase(ACAT). The results showed that the mixed nitrogen source is slightly better than the organic nitrogen source, and it is more beneficial to β-oxidation cycle (Fig. 5A and B).

In the analysis of microbial diversity, bacteria *Psychrilyobacter* are one of the dominant bacteria groups. And *Psychrilyobacter* can degrade nitramine explosives at low temperature conditions(40). So we analyzed the copy number of the genes encoding nitric oxide synthase(*xplA*) (Fig. 5B). Reductive dehalogenases are responsible for biological dehalogenation in organohalide-respiring bacteria, with substrates including polychlorinated biphenyls or dioxins, which are usually membrane associated and oxygen sensitive(18, 29). So in view of the high chloride ion in the marine environment, we also analyzed the copy number of the genes encoding reductive dehalogenase (VcrA_Vcr2).

The results showed that organic and inorganic nitrogen mixture can enhance the copies of *xplA* gene and *vcrA_vcr2*, and promote the degradation of nitrogen and chlorinated compounds in oil (Fig. 5B).

The effects of different nitrogen sources on microbial community structure, biodegradation functional genes and oil degradation rates were discussed in this research. Addition nitrogen source can promote oil biodegradation, but the effect of adding organic nitrogen sources or mixture nitrogen sources on oil degradation generally better than that of inorganic nitrogen sources, which indicated that organic nitrogen source was better than inorganic nitrogen source on oil biodegradation; and for nitrogen source mixture, the proportion of different nitrogen sources was important for oil biodegradation. The type of nitrogen source and their proportion all affect the oil biodegradation effect, and have also obviously influence on microbial structure and biodegradation functional gene distribution. We should pay attention to the reasonable collocation of organic nitrogen source and inorganic nitrogen source to improve the rate of oil biodegradation.

## Discussion

The metabolic process is closely related to the microbial communities in the surrounding environments, so, distribution characteristics of microbes with different nitrogen sources is also important for improving the effect of petroleum degradation by using biostimulation methods. Most biostimulation methods use inorganic nitrogen sources as the main nutritional supplement to improve the efficiency of petroleum degradation. Louati et al. (15) reported that the biodegradation rate of phenanthrene was up to 98% after adding nitrogen fertilizer or mineral salt medium. Similar results have been obtained in the removal of anthracene in the Bizerta lagoon sediments and oil spill remediation by using particle inorganic fertilizers (26, 28). The experimental results in this research showed that the effects of different nitrogen sources on microbial oil degradation were obviously different, and the promotion effect of organic nitrogen sources was obviously better than that of inorganic nitrogen sources, and the promotion effect of the optimized nitrogen combination was better than that of organic nitrogen source.

When oil pollution incident occurred, the leaking oil could be used as carbon and energy sources to change the structure of microbial communities; at the same time, microorganisms which could consume oil were rapidly and strongly selected (2, 32). Previous study showed that *Proteobacteria* was ubiquitous in the contaminated environment, which was an important oil degrader at aerobic culture condition (9, 10). In this research, the microbial diversity of marine sediments enriched with different nitrogen sources decreased with the increase of oil degradation rate, and the population of microorganisms migrated obviously when enriched with different nitrogen sources. *Proteobacteria* was still the main oil degrading bacteria at anaerobic condition, but the reasonable proportions of *Proteobacteria, Bacteroidetes* and *Fusobacteria* made the greatest contribution to petroleum degradation. At the same time, the results of the analysis of oil degradation rate showed that oil biodegradation rates in samples enriched with organic nitrogen sources were higher than that in the samples with inorganic nitrogen sources, and the highest oil biodegradation rate was in sample YH. In fact, since the lack of nutrition at deep sea floor, the rate-limiting process in bioremediation of oil pollution *in situ* is the provision of nutrition. The result in this research showed that supplement of nitrogen was one of the efficient oil biodegradation methods (5, 22). Thus, the nitrogen requirement in deep seafloor can be solved by addition nitrogen fertilizers, such as ammonium, nitrates and urea. According to the results above, it is more important to supplement the reasonable allocation of nitrogen source, obviously.

The microbial composition in YH samples with the highest oil degradation rate was analyzed. the 5 genera such as *Marinifilum, Psychrilyobacter, Propionigenium, Psychromonas*, and *Vibrio* showed that the highest relative abundance. *Marinifilum* and *Psychromonas* were facultative anaerobic bacteria. Wang et al. reported that *Alcanivorax, Marinobacter, Novosphingobium, Rhodococcus* and *Pseudoalteromonas* were found to be predominant oil-degrading bacteria in the polluted seawater and sediments from Bohai Bay and Yellow Sea (34, 36). Obviously, the microbial population at low temperature anaerobic condition revealed in this study was greatly different from that reported by Wang et al. *Marinifilum (*35.7%) and *Psychrilyobacter* (26.6%) were the dominant bacteria groups in YH samples, few reports about *Marinifilum and Psychrilyobacter* can degrade oil. Most strains of the genus *Marinifilum* were isolated from marine environments,which is also the group of living bacteria in the intestines of some marine organisms (21), but the role of *Marinifilum* that plays in marine ecology and the life cycle of marine organisms is not yet known. Our experimental results showed that *Marinifilum* with the highest relative abundance of microorganisms may play important role in the process of petroleum degradation, which may be an potential bacteria with high degradation oil ability. *Psychrilyobacter*, was obligately anaerobic marine bacteria, can degrade nitramine explosives at low temperature conditions (40). XplA (Nitric oxide synthase) is detected only in explosives contaminated sites thereby suggesting rapid catabolic activity to be carried out by this enzyme on RDX (hexahydro-1,3,5-trinitro-1,3,5-triazine) (37). And XplA was mainly responsible for initiating cyclic nitroamines degradation, the explosive RDX degradation, involving sequential reduction of N-NO_2_ to the corresponding N-NO groups at anaerobic condition (4). The results showed that organic and inorganic nitrogen mixture can enhance the copies of *xplA* gene, and promote the degradation of petroleum (Fig. 5B). Few reports about *Propionigenium* and *Psychromona* species can degrade oil, but *Vibrio* species were reported to be able to degrade oil (38). The results imply that the combined nitrogen source can be used to enrich and screen anaerobic oil degrading microorganisms effectively.

Through the analysis of the key functional genes in the anaerobic oil degradation pathway, the molecular mechanism of the best combination nitrogen source to promote oil degradation was revealed. The main pathway of the anaerobic metabolism of alkanes is the addition reaction of fumarate. This reaction is first carried out by Alkylsuccinate synthase (ASS) or Benzylsuucinate synthase (BSS) (31). Alkane activation is achieved via conjugation with fumarate at C-2 position by the alkyl succinate synthase (ASS)/ Benzylsuucinate synthase (BSS) to yield (1-methyl-alkyl/Benzyl) succinate. After addition of a coenzyme A (CoA) to the product via the action of succinyl-CoA/benzylsuccinate CoA-transferase SAcT and carbon rearrangement and decarboxylation catalyzed by acelyl-CoA carboxylase carboxyl transferase (ACCT), a methylated fatty acid that is two carbons larger than the original n-alkane is formed (3). The resulting fatty acid is funneled into β-oxidation pathway (Fig. 5A). This reaction is the universally recognized anaerobic degradation of petroleum by many anaerobic bacteria, including denitrifying microorganisms, sulphate-reducing bacteria, methanogenic consortia and metal-reducing (Mn(IV), Fe(III)) bacteria (1). Metagenomic analysis illuminated the mixed nitrogen source promoted the transfer of succinyl-CoA, synthesis of acetyl CoA and β-oxidation cycle, and then was beneficial to degradation of petroleum relatively at low temperature anaerobic condition.

## Acknowledgments

This work was supported by the National Natural Science Foundation of China (No.21276047, No.31600004 and No.31770006), the Fundamental Research Funds for the Central Universities (wd01187, wd01189).

